# Asian lineage of Zika virus RNA pseudoknot may induce ribosomal frameshift and produce a new neuroinvasive protein ZIKV-NS1’

**DOI:** 10.1101/105809

**Authors:** Tiago Tambonis, Vinícius G. Contessoto, Cíntia Bittar, Marília F. Calmon, Maurício L. Nogueira, Paula Rahal, Vitor B. P. Leite

**Affiliations:** São Paulo State University (UNESP), Department of Physics, São José do Rio Preto - SP, 15054-000, Brazil; Brazilian Bioethanol Science and Technology Laboratory (CTBE), Campinas - SP, 13083-100, Brazil; São Paulo State University (UNESP), Department of Biology - Genomic Studies Laboratory, São José do Rio Preto - SP, 15054-000, Brazil; São José do Rio Preto School of Medicine (FAMERP), São José do Rio Preto - SP, 15090-000, Brazil

**Author notes:** These authors contributed equally to this work (Tiago Tambonis, Vinícius G. Contessoto).

## Abstract

Zika virus (ZIKV) is a threat to humanity, and understanding its neuroinvasiveness is a major challenge. Microcephaly observed in neonates in Brazil is associated with ZIKV that belongs to the Asian lineage. What distinguishes the neuroinvasiveness between the RNA lineages from Asia and Africa is still unknown. Here we identify an aspect that may explain the different behavior between the two lineages. The distinction between the two groups is the occurrence of an alternative protein NS1’ (ZIKV-NS1’), which happens through a pseudoknot in the virus RNA that induces a ribosomal frameshift. Presence of NS1’ protein is also observed in other *Flavivirus* that are neuroinvasive, and when NS1’ production is suppressed, neuroinvasiveness is reduced.^1^ This evidence gives grounds to suggest that the ZIKV-NS1’ occurring in the Asian lineage is responsible for neuro-tropism, which causes the neuro-pathologies associated with ZIKV infection, of which microcephaly is the most devastating. The existence of ZIKV-NS1’, which only exists in the Asian lineage, was inferred through bioinformatic methods, and it has yet to be experimentally observed. If its occurrence is confirmed, it will be a potential target in fighting the neuro-diseases associated with ZIKV.

---

Zika virus (ZIKV) is an *Arbovirus* belonging to the Flaviviridae family and *Flavivirus* genus like Dengue virus (DENV), West of Nile virus (WNV), Yellow Fever virus (YFV) and Japanese encephalitis virus (JEV). The ZIKV genome is similar to other viruses in the Flavivirus genus and encodes three structural proteins and seven non-structural (NS) proteins.^2^ ZIKV was first isolated in Africa,^3^ and subsequently an Asian lineage was also reported.^4, 5^ Neurological complications such as Guillain-Barré syndrome in adults in French Polynesia and microcephaly in newborns in Brazil were associated with the Asian ZIKV lineage.^6–8^ There is well-known evidence of neurological problems associated with *Flavivirus* genus.^9^ In particular, for JEV and WNV, the production of an eleventh protein related to the viral neuroinvasiveness was observed, the NS1’.^1, 10^ This protein was first observed in the 80s in JEV-infected cells,^11^ and its production mechanism was recently discovered.^10^ NS1’ is produced as the result of a −1 ribosomal frameshift at the 5’ end of the NS2A coding sequence, which consists of the NS1 protein with the addition of 52 more residues.^12^ The −1 ribosomal frameshift occurs through a pseudoknot formation preceded by a ‘slippery’ heptanucleotide, which is followed by a ‘spacer’ region.^13^ This mechanism is also observed in other viruses.^14^

Here, using bioinformatic approaches, we investigated the likelihood of a ribosomal frameshift occurrence in ZIKV, which may account for the translation of a NS1’ protein. 29 different Zika virus gene sequences from GenBank^15^ were analyzed, of which 15 were from Asian lineage and 14 from African lineage. Probing for the conditions for the occurrence of a ribosomal frameshift, a ‘slippery’ heptanucleotide CCCAAAU and a pseudoknot formation propensity in a position near the extremity of the NS1 gene of ZIKV was observed. These features allow the ribosomal frameshift to happen in ZIKV, in similar positions to those occurring in JEV and WNV. This frameshift occurrence in ZIKV was observed in all Asian lineage sequences but not in African lineage sequences. The Brazil strain in which microcephaly cases were observed comes from the Asian lineage.^16, 17^ On the other hand, no cases of microcephaly associated with the ZIKV African lineage were reported.^18^

The predicted −1 frameshift in the ZIKV RNA code is located at nucleotide position 794 in the NS1 region, allowing a translation of 40 different residues until the stop codon *UAA* appears at nucleotide position 913 after the occurrence of the frameshift. The translated ZIKV-NS1’ protein has 303 residues with~35 kDa of molecular weight. ZIKV-NS1’ is 49 residues smaller than the NS1 protein, and both proteins are similar in residue sequence until position 263, but after this position the residue sequences are completely different. The ZIKV-NS1’ amino acid sequence based on the KU321639 Genbank entry is: GCSVDFSKKETRCGTGVFVYNDVEAWRDRYKYHPDSPRRLAAAVKQAWEDGICGISSVSRMENIMWRSVEGELNAILEENGVQLTVVVGSVKNPMWRGPQRLPVPVNELPHGWKAWGKSHFVRAAKTNNSFVVDGDTLKECPLKHRAWNSFLVEDHGFGVFHTSVWLKVREDYSLECDPAVIGTAVKGKEAVHSDLGYWIESEKNDTWRLKRAHLIEMKTCEWPKSHTLWTDGIEESDLIIPKSLAGPLSHHNTREGYRTQMK**ARGGNMWNKRTISEINHCKRKGDRGMVLQGVHNAPTVVPG**, in which the bold part of the sequence is associated with the 40 residues that are translated after the pseudoknot, and they are nearly identical (two different AA at the most) for all ZIKV Asian sequences. The results, along with experimental evidence from JEV and WNV studies,^1, 10^ suggest that the Asian lineage ZIKV RNA has the essential characteristics to produce the pseudoknot, which allows for a ribosomal frameshift to produce the unexpected ZIKV-NS1’ protein. Moreover, the appearance of ZIKV-NS1’ protein may be associated with the neuroinvasiveness of ZIKV, and may be related to microcephaly in neonates.

Studies in JEV and WNV show that the addition of mutations in nucleotide which makes it difficult for the pseudoknot in viral RNA to form, correlates with the reduction of neuroinvasiveness in mice.^1, 19, 20^ It is plausible to assume that ZIKV neuroinvasiveness may be related to the occurrence of ZIKV-NS1’, just like to the NS1’ observed in JEV and WNV. ZIKV-NS1’ is expected to be rarely expressed, similarly to NS1’ for JEV and WNV, but should be experimentally detectable in a straightforward manner.

A recent study of ZIKV with x-ray crystallography showed pseudoknot formation in its RNA, and these structures were associated with exonuclease Xrn1 resistance, which also points to the importance of fully understanding pseudoknot formation and function.^21^ Experimental groups with the necessary expertise are urged to validate this conjecture. The corroboration of the existence of ZIKV-NS1’ protein may open a window of opportunity in the understanding of its relationship with microcephaly and ZIKV neuroinvasiveness, as well as in the development of drugs and vaccines with this specific target.

## Methods

The frameshift mechanism consists of a ‘slippery’ heptanucleotide followed by a ‘spacer’ region of 5-9 nucleotides and then a stable RNA secondary structure such as a pseudoknot. The ‘slippery’ heptanucleotide consensus sequence is NNNWWWH, where NNN are any three identical nucleotides, WWW represents AAA or UUU and H represents A, C or U.^14^ The 29 studied gene sequences of Zika virus were obtained from GenBank,^15^ being 15 from Asian lineage: KU497555, KU729218, KU321639, KU729217, KU365778, KU365777, KU365780, KU365779, KU707826, KJ776791, KU922923, KU922960, KU820899, KU744693, KU963796 and 14 from the African lineage: KF268948, KF268950, KF268949, AY632535, LC002520, HQ234501, HQ234500, KF383117, NC 012532, KF383118, KF383115, KF383119, DQ859059, KF383121. All possible ‘slippery’ heptanucleotide combinations were exhausted constructed and performed in R environment^22^ *version 3.3.2*. The formation of pseudoknot structures was analyzed using *pKiss* tool ^23^ from nucleotide sequence: GGCCAUGGCACAGUGAAGAGCUUGAAAUUCGGUUUGAGGAAUGCCCAGGCACUAAGGUC and structure [[[[............((((((((.((((........))))...)))))))).]]]]. All sequence alignments were executed using the MUSCLE algorithm^24^ from *Bioinformatic toolkit* web server.^25^

## Acknowledgements

TT was supported by Coordination for the Improvement of Higher Education Personnel (CAPES), Brazil. VGC was funded by grant 2016/13998-8, São Paulo Research Foundation (FAPESP)and CAPES. VBPL was supported by the National Council for Scientific and Technological Development (CNPq) and FAPESP grant 2014/06862-7. CB was supported by the grant CNPq 165802/2015-4. MFC, MLN and PH supported by FAPESP grant 2014/22198-0 and by CNPq grant 440723/2016-7.

### Author Contributions

C.B., P.R., T.T., V.G.C. and V.B.P.L. conceived the experiment, T.T. and V.G.C. conducted the experiment and analysed the results. All authors reviewed the manuscript.

### Author Information

Correspondence and requests for materials should be addressed to V.B.P.L.

